# End resection and telomere healing of DNA double-strand breaks during nematode programmed DNA elimination

**DOI:** 10.1101/2024.03.15.585292

**Authors:** Brandon Estrem, Richard E. Davis, Jianbin Wang

## Abstract

Most DNA double-strand breaks (DSBs) are harmful to genome integrity. However, some forms of DSBs are essential to biological processes, such as meiotic recombination and V(D)J recombination. DSBs are also required for programmed DNA elimination (PDE) in ciliates and nematodes. In nematodes, the DSBs are healed with telomere addition. While telomere addition sites have been well-characterized, little is known regarding the DSBs that fragment nematode chromosomes. Here, we used embryos from the nematode *Ascaris* to study the timing of PDE breaks and examine the DSBs and their end processing. Using END-seq, we characterize the DSB ends and demonstrate that DNA breaks are introduced before mitosis, followed by extensive end resection. The resection profile is unique for each break site, and the resection generates 3’ overhangs before the addition of telomeres. Interestingly, telomere healing occurs much more frequently on retained DSB ends than on eliminated ends. This biased repair of the DSB ends in *Ascaris* may be due to the sequestration of the eliminated DNA into micronuclei, preventing their ends from telomere healing. Additional DNA breaks occur within the eliminated DNA in both *Ascaris* and *Parascaris*, ensuring chromosomal breakage and providing a fail-safe mechanism for nematode PDE.

## INTRODUCTION

Programmed DNA elimination (PDE) is an exception to the paradigm of genome integrity (1–4). It removes DNA from the germline genome to generate a reduced somatic genome within the life cycle of an organism. PDE occurs in single-cell ciliates (5–7) and a growing list of metazoans (8–11), suggesting the PDE process likely evolved independently in distinct phylogenetic groups and confers biological function(s). There are significant variations in how PDE occurs among diverse organisms, including the developmental stages where it occurs, the amount and types of DNA eliminated, and the genomic consequences of PDE (1–4). However, the overall functions of PDE in metazoans remain speculative as an experimental model where PDE is fully blocked has yet to be established.

Two distinct mechanisms are used to eliminate DNA during PDE. In the first, the entire chromosome(s) is lost, likely through heterochromatinization and asymmetric division or the loss of lagging chromosomes (8,9). Loss of entire chromosome(s) occurs in some arthropods, birds, lampreys, hagfish, and mammals. In the second mechanism, chromosomes are broken, and specific fragments are reproducibly retained or lost (10,12). Chromosome fragmentation requires the generation of DNA double-strand breaks (DSBs) and their subsequent healing. PDE-associated DSBs have been identified in ciliates and some nematodes; they are likely also present in some copepods (13,14) and hagfish (15,16).

The generation and healing of DSBs differ between ciliates and nematodes. In most ciliates, two types of genome changes occur that require DSBs. The majority of the DSBs are generated during the excision of internal eliminated sequences (IESs) by domesticated transposases, followed by the fusion of broken ends through non-homologous end joining (NHEJ) mediated repair (5–7). The second form of DSBs occurs during chromosome fragmentation at the chromosome breakage sequences (12). This process is coupled with *de novo* telomere addition using telomerase (17,18). In comparison, much less is known about the molecular mechanism of DSBs in nematodes. The DSBs occur at the ends of all nematode chromosomes (19,20) as well as in the middle of some chromosomes (21). The DSBs are healed by *de novo* telomere addition and become the ends of new somatic chromosomes (22–25). Thus, PDE in nematodes removes and remodels the ends of all germline chromosomes and generates new somatic chromosomes.

In nematodes, telomere addition sites have been characterized using genome sequencing (20,24). In the human and pig parasitic nematode *Ascaris* (26), we previously identified 72 chromosomal breakage regions (CBRs) where telomeres are added. These CBRs occupy a 3-6 kb window (24,27,28) and are not associated with specific sequence motifs, common histone marks, or small RNAs, suggesting the sites for the telomere addition are sequence independent. However, all CBRs are associated with more accessible chromatin during DNA elimination, indicating specific mechanisms are involved in identifying the sites for chromosomal breakage and telomere addition (24). In contrast, *de novo* telomere addition sites in the free-living nematode *Oscheius tipulae* reside primarily at a discrete site in the center of a 30-nt palindromic motif called SFE (<u>S</u>equence <u>F</u>or <u>E</u>limination) (20). Our initial END-seq analysis indicated the DSBs in *O. tipulae* were resected to generate long 3’-overhangs, and telomeres were unbiasedly added to both the retained and eliminated ends of the DSBs (20). However, due to the fast cell cycle (20-30 minutes/cycle) in *O. tipulae*, we could not determine the timing of DSBs or the dynamics of DNA end resection and telomere addition. Furthermore, the potential molecular differences of the DSBs, end processing, and telomere addition between a motif-based (SFEs in *O. tipulae*) and a sequence-independent (CBRs in *Ascaris*) process remain largely unknown.

Here, we determine the timing, nature, and sequence features associated with the DSBs for *Ascaris* PDE. Using synchronized embryos, we carried out END-seq to characterize DSBs and end resection at specific stages of the cell cycle during PDE (4-8-cell embryos). These stages cover discrete time points of *Ascaris* PDE, including the onset of DSBs, end resection, new telomere addition, and DNA degradation. Our data demonstrates that the DSBs are introduced at the G2 phase before mitosis and are followed by extensive end resection. The DSBs occur heterogeneously within the CBRs. Moreover, telomeres are mainly added to the retained ends of DSBs in *Ascaris*, while the eliminated ends undergo further resection – in contrast to the unbiased telomere healing observed in *O. tipulae* (20). We also identified additional DSBs within the eliminated DNA in *Ascaris* and the related horse parasitic nematode *Parascaris*. In combination with the alternative breaks in *O. tipulae* (20), these additional DSBs suggest a common fail-safe mechanism, where additional DNA breaks occur to ensure PDE in these nematodes. Furthermore, telomere healing of DSBs appears to be a specific process associated with PDE, as exogenously introduced DSBs are not healed by telomere addition. Overall, our results provide insights into the DSBs and telomere healing and reveal variations in the molecular processes of PDE in diverse nematodes.

## MATERIALS AND METHODS

### Sample collection and embryo development

*Ascaris* females were collected, and the fertilized embryos (0hr, 1-cell before prenuclear fusion) were harvested and processed as previously described (29,30). *Ascaris* 0hr samples were incubated at 30°C with constant shaking for the desired time (from 50hr to 98hr; see Fig. 1 for the population average of cell stage and phase of cell cycle). For all molecular experiments, the chitinous eggshells were first digested with base-bleach treatment (0.4 M KOH, 2% sodium hypochlorite [Fisher Scientific, catalog #SS290-1]) for 1.5 hours at 30°C. *Parascaris* samples were collected as described (21,24). *Parascaris* eggs were prepared similarly to *Ascaris*, except the incubation was carried out at 37°C, and the embryonation time was shorter (10-14hr). The *Parascaris* embryos used for the END-seq library were from mixed stages of 1-2 cells (before PDE) and 2-8 cells (during PDE).

**Figure 1.**
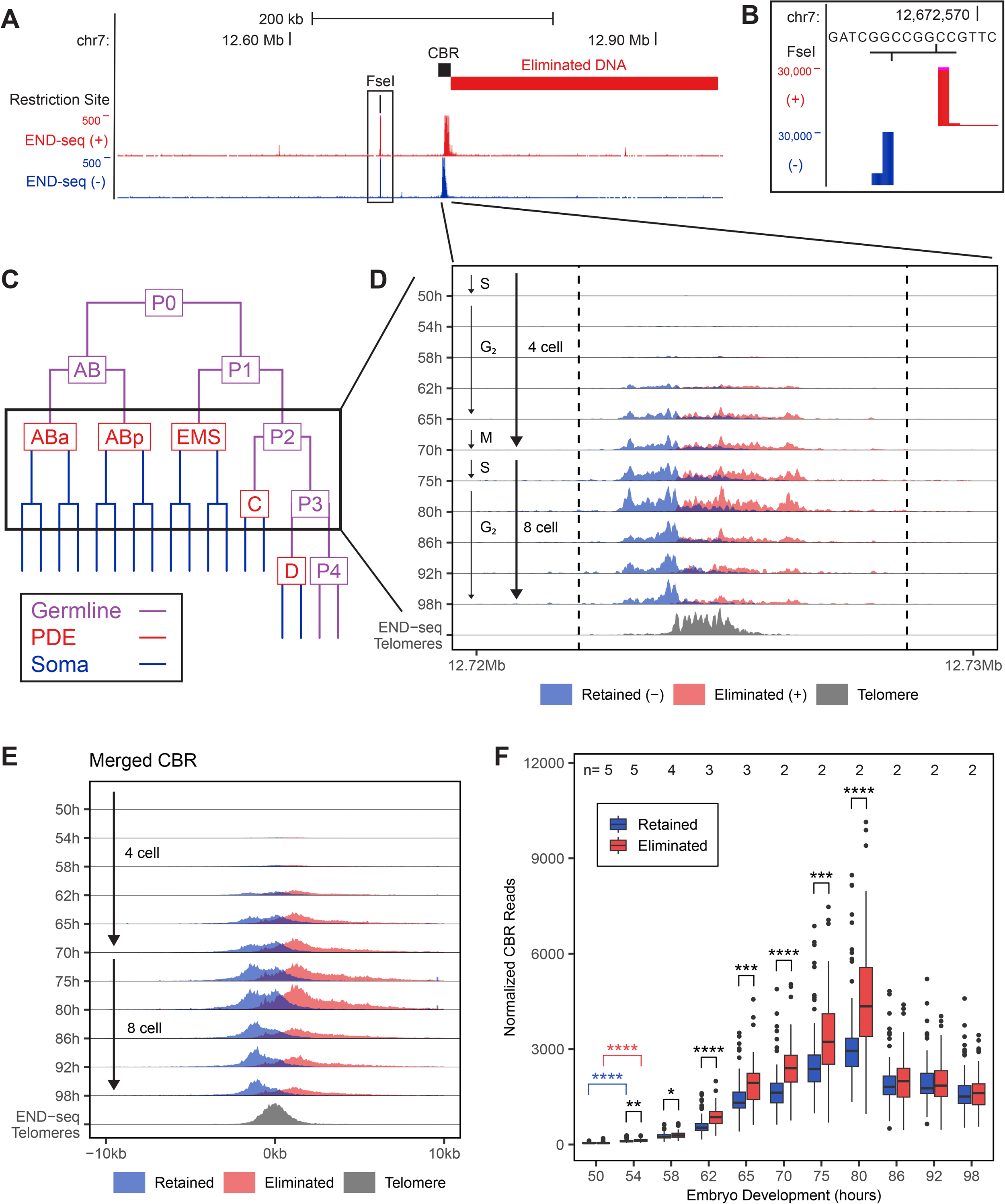
*Ascaris* DSBs for PDE are generated before mitosis. **A)** END-seq identifies DSBs in *Ascaris* embryos. A Genome Browser view of END-seq reads from 70hr (4-6-cell) embryos. Shown is a 500 kb region containing an FseI restriction site and a CBR. Reads were split by strand and colored red (+) and blue (−). **B)** Inset of the FseI site boxed in **A**. FseI generates a 4-nt 3’-overhang that is blunted during END-seq, as indicated by the high number of END-seq reads matching the blunted ends, leaving a 4-nt gap in END-seq signal between the two strands. **C)** *Ascaris* cell lineage during early development. Germ cells are purple, cells that undergo PDE are red, and somatic cells are blue. **D)** Timing of DSBs detected by END-seq. Ridgeline plot of normalized END-seq reads across 11 developmental stages (y-axis) at the same CBR as in **A** and its flanking regions (x-axis; total 14 kb with 100 bp bins and 10 bp sliding window). Dashed lines mark the boundary of END-seq signal enrichment (defined with MACS3). Reads are colored by strand (red and blue), and telomeric reads are grey. Estimates of cell number and phase of the cell cycle from the population of embryos are indicated on the left. **E)** Average END-seq profile across all CBRs. The 72 CBRs were aligned by the median telomere addition sites at each CBR and the END-seq coverage was merged to create an average profile. Legend same as in **D**. **F)** A bias of resection in the retained vs. eliminated DSBs ends. The average END-seq read counts at each CBR were plotted for each developmental stage. The number of END-seq libraries (biological replicates) for each stage is indicated at the top of the graph. Many time points (from 54hr to 80hr) have more END-seq reads in the eliminated sides than the retained ends. All time points have significantly more END-seq reads than 50hr (significance only shown between 50hr and 54hr). Wilcoxon statistic tests were used with * p < 0.05, ** p < 0.01, *** p < 0.001, **** p < 1e-4.

### END-seq library preparation

The de-coated embryos were treated with 90% isopropanol for one minute to remove the outer membrane, followed by 3X washes in PBS before proceeding with END-seq procedures (31,32). Briefly, embryos were embedded in agarose plugs to protect the DNA from exogenous breaks. For each plug, we used ∼50 uL of packed embryos as the starting material. Some plugs were treated with the restriction enzymes AsiSI, FseI, AscI, and/or PmeI (NEB, catalog #’s R0630, R0588, R0558, and R0560) to generate DSBs as internal controls. DSBs were blunted with exonuclease VII (NEB, catalog # R0630) and exonuclease T (NEB, catalog # M0625). Blunt ends were A-tailed and capped with END-seq adaptor 1, a biotinylated hairpin adaptor (32). Plugs were melted at 70°C and treated with β-Agarase I (NEB, catalog #M0392) to liberate the DNA. The DNA was then sheared to 200-300 bp with a Covaris M220 focused ultrasonicator (130 uL tube [Covaris part number 520045], 4°C, peak power 50, duty 16, cycles/burst 200 for 420 seconds). DNA fragments containing END-seq adaptor 1 were isolated with Dynabeads MyOne Streptavidin C1 (Invitrogen, catalog #65001). END-seq adaptor 2 was ligated to the sheared ends of the A-tailed DNA fragments. The hairpins within the adaptors were digested with USER (NEB, catalog #M5505), and the DNA was amplified with Illumina TruSeq primers and barcodes. The libraries were sequenced with Illumina HiSeq 2500 or NovaSeq 6000 at the University of Colorado Anschutz Medical Campus Genomics Core.

The following modifications were made to the END-seq protocol to capture DNA with blunt ends and/or 5’- and 3’-overhangs. For the direct capture method, we excluded the exonuclease VII and exonuclease T treatments to only capture blunt ends. For the all-END protocol, the plugs were treated with T4 polymerase (with dNTPs) to allow the filling of overhangs before exonuclease treatment, thus capturing DNA ends with blunt, 5’- and 3’-overhangs. These experiments were done on 68hr embryos when endogenous DSBs from PDE are abundant. The samples were also treated with restriction enzymes AsiSI (3’-overhang, 3,385 sites in the genome), AscI (5’-overhang, 416 sites), and PmeI (blunt, 5,588 sites) to generate control DSBs.

### Southern blotting

High-molecular-weight DNA was extracted from the germline (ovary), four stages of early embryos (50-74hr), and somatic cells (7-day embryos) using an agarose embedding method (27). About 3 µg of DNA from each sample was digested with two restriction enzymes (PstI; NEB catalog #R0140 and XhoI; NEB catalog #R0146). The digested genomic DNA was resolved on a 1%, 0.5X TBE buffer agarose gel with a 1 kb Plus DNA Ladder (Invitrogen, catalog #12308-011). The DNA was transferred to a Hybond-XL membrane using 0.5 N NaOH/1.5 M NaCl. The membrane was treated with 1200 µJoules in a UVP CL-1000 Crosslinker. We selected a 700 bp region within the retained side of a CBR (CBR_m6b, see Fig. 3E) as the probe for hybridization. The PCR amplicon (primers: forward = TTTCTAAGACTCTCTCCCGTA and reverse = GATTAGAAGTAGCCGACCAA) was labeled with dCTP [α-^32^P] using Random Primer DNA Labeling Kit Ver.2.0 (Takara, catalog #6045). The hybridization was done at 65°C overnight in Church and Gilbert Moderate Hybridization Buffer (1% BSA, 500 mM sodium phosphate, 15% formamide, 1 mM EDTA, and 7% SDS) and washed in 0.2x SSC, 0.1% SDS at 55**°**C using a GENE Mate HO6000V hybridization oven. The blot was imaged using an Amersham Typhoon Biomolecular Imager.

### X-ray irradiation

*Ascaris* embryos (65hr [4-cell] or 70hr [4-6-cell] with eggshell removed) were placed in 60 mm petri dishes and irradiated with 100 or 200 Gy of X-rays in an RS 2000 small animal irradiator (∼4 Gy/min at shelf level 5). Control samples were placed in 60 mm petri dishes and left on the counter for the same period while the X-ray sample was irradiated. To determine the impact of X-ray irradiation on *Ascaris* embryo development, we allowed treated embryos to recover for 24hr or 48hr at 30°C post-irradiation (corresponding to 1-3 cell cycles). After recovery, the number of cells in the embryos was counted using light microscopy and Hoechst staining. For the staining, ∼120 uL of packed embryos were treated with 90% isopropanol for 1 minute, washed in PBS pH 7, and were subjected to the stain using Hoechst 33342 (1 mg/mL) (Invitrogen, Fisher Cat# H3570) following the procedures as described previously (20).

For END-seq, the irradiated embryos were embedded in agarose plugs and processed as described above. For genomic DNA isolation, ∼80 uL packed irradiated embryos and control embryos were resuspended in 2 mL buffer G2 (Qiagen, catalog # 1014636) and digested with proteinase K (1 mg/mL) (Invitrogen, catalog #AM2544). The embryos were lysed with five strokes in a 7-mL metal dounce followed by a 2hr incubation at 37°C. The lysate was centrifuged 5,000 x g for 10 minutes to pellet debris. The supernatant was processed with Genomic-tip 20/G columns (Qiagen catalog #10223) to prepare genomic DNA. Genomic libraries were made using Illumina DNA Preparation Kit (Cat# 20018704) and sequenced with Illumina NovaSeq 6000.

### END-seq data mapping and visualization

For all END-seq analyses, only the read 1 (R1) file with the captured DSB ends was processed. END-seq reads containing two consecutive telomeric repeat units (TTAGGCTTAGGC or the reverse complement GCCTAAGCCTAA) were first identified from sequencing files using an in-house Perl script; they were filtered and used for the analysis of *de novo* telomere addition (see below). The rest of the END-seq reads were mapped to the appropriate reference genome (*Ascaris* v3 [accession #: JACCHR000000000] or *Parascaris* v2 [accession #pending]) with bowtie2 (local alignment) (33) and processed with SAMtools (34) to generated bam files. For samples treated with restriction digestion, BEDTools (35) intersect was used to remove reads mapped to restriction sites. The 5’-end position of each read was mapped, separated by strand, and normalized to ten million genome-mapped reads using BEDtools *genomecov* (35). The mapping results were converted to bigWig format using bedGraphToBigWig and loaded into UCSC genome browser track data hubs (36).

### Identification of *de novo* telomere addition

To analyze new telomere addition during PDE developmental stages, END-seq reads containing two consecutive telomere repeats were converted to the G-rich strand (TTAGGC). To identify reads with non-telomeric sequences, we first mapped the full length of these reads (without trimming or clipping) to the germline genome using bowtie2 end-to-end alignment (33). Those mapped reads were false-positive telomeric reads and were removed from the downstream analysis. The rest of the reads were trimmed with fastx_clipper (http://hannonlab.cshl.edu/fastx_toolkit/index.html) using “-v -n -l 25 -a TTAGGCTTAGGC” parameters. The trimmed reads were mapped to the genome with bowtie2 (local alignment), and the mapping results (bam files) were processed as described above, except the 3’-positions (the sites where new telomeres are added) rather than 5’-positions were obtained with BEDTools *genomecov*. To filter ambiguous mapping results, reads with < 50 bp of genomic sequence were excluded from the analysis since few of them have >25 bp of unique sequence after removing the telomeric portion of the reads.

### Identification of break sites and resection boundaries

To identify genomic regions with enriched END-seq signal, representative libraries from each stage of PDE and a control library (before PDE) were first split by forward (+) and reverse (−) strands. Each strand was independently analyzed with MACS3 (37) *callpeak* (*Ascaris*: -g 2.43e8 -s 120 --nomodel --broad --min-length 1000; *Parascaris:* -g 2.40e8 --nomodel –broad --broad-cutoff 0.13). The MACS3 output was filtered to remove peaks only found on one strand (BEDTools window: *Ascaris* -w 1000 and *Parascaris* -w 2000). In *Parascaris*, due to the limited END-seq signal, some CBRs and their resection boundaries were defined manually by further assessing genome read coverage and telomere addition sites using the genome browser.

To identify alternative break sites, MACS3 peaks in the eliminated regions were assessed. Peaks overlapping with highly repetitive regions were removed from downstream analysis. END-seq reads were first mapped to the CBRs to ensure the reads in the eliminated regions were not derived from the existing 72 CBRs due to their potential of being repetitive sequences to the CBRs; then, the remaining reads were mapped to the rest of the genome using bowtie2. After the sequential mapping, non-CBR regions with a significant number of reads were considered as alternative CBRs. To identify if alternative CBRs could have repetitive sequences similar to the 72 CBRs after the sequential mapping, the overall coverage of the END-seq was used to determine if multiple CBRs with expected multiple-fold read coverage exist, as demonstrated by the PDE breaks in the nematode *O. tipulae* (20).

The END-seq signal region (defined by MACS3) for each CBR was extended to 20 kb to include a flanking region for comparative analyses of all CBRs across developmental stages. The 20 kb region was binned into 100 bp windows with a sliding window of 10 bp using BEDTools *makewindows* (-w 100 -s 10). Normalized END-seq data for each stage of development was merged with BEDTools *unionbedg* and mapped to the binned 20 kb break regions using BEDTools *map* (-c 4 -o mean -null 0). END-seq *de novo* telomere data was independently normalized, merged, and processed using the same approach. The data was plotted using R using packages *tidyverse*, *reader*, *scales*, *ggpubr*, *ggridges*, and *extrafont*.

For meta-analysis of all CBRs (such as in Fig. 1E), the CBRs were aligned by the median END-seq telomere read. A 20 kb region centered on the telomere median was binned with BEDTools *makewindows* (-w 100 -s 10). The coordinates were converted to a relative scale from −10,000 to 10,000 bp, and CBRs with eliminated DNA on the left were inverted, so all CBRs were in the same orientation (eliminated DNA on the right). The same process was used to generate the merged *Parascaris* plot, except the median break site was determined using telomere addition sites from the somatic tissue (24).

### Simulation of END-seq profiles using the *O. tipulae* resection profile

The overall END-seq resection profile from 12 canonical break sites was obtained from *O. tipulae* (20). The 5’ read coordinates were converted to the relative distances from the telomere addition site, normalized to the number of telomeres, and oriented so that eliminated reads were on the right-hand side. This pattern was used to simulate an END-seq profile at each *Ascaris* CBR, assuming a DSB gives rise to a similar resection profile. The *O. tipulae* END-seq pattern was applied to the positions and frequencies of the observed telomere addition sites from the wild population of *Ascaris* embryos. The simulated profiles were compared with observed END-seq profiles in ridgeline plots (see Fig. 2E and Fig. S2).

**Figure 2.**
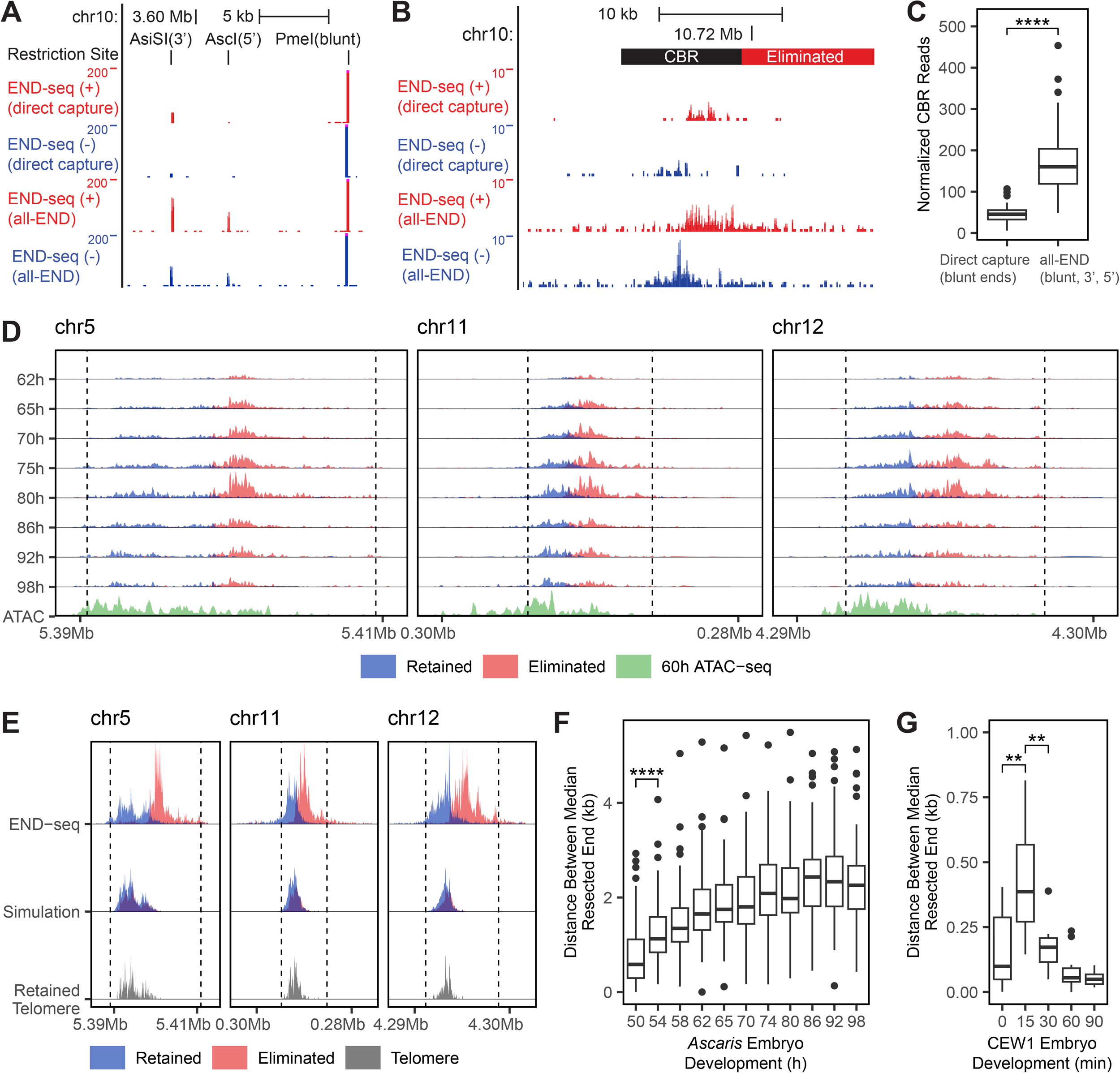
*Ascaris* DSBs undergo extensive end resection. **A)** Modifications of the END-seq procedure capture DSBs with different end features. The direct capture method excluded the exonuclease treatments and thus only identified DSBs with blunt ends; while the all-END protocol treated the samples with T4 DNA Polymerase to generate blunt ends from both 5’- and 3’-overhangs for their subsequent capture (see Methods). Shown is a Genome Browser view of END-seq reads from 68hr embryos treated with AsiSI (3’-overhang), AscI (5’-overhang), and PmeI (blunt) restriction enzymes. **B)** Most of the *Ascaris* DSB ends have an overhang structure at their ends. Shown is an exemplary CBR region from direct capture and all-END experiments. Libraries were normalized to the same number of mapped reads. **C)** Quantification of normalized reads in each CBR from direct capture and all-END experiments. **D)** Each CBR has a distinct resection profile that may be influenced by the local sequence, nucleosome organization, and chromatin structure. Three exemplary CBRs (17 kb) with END-seq and ATAC-seq data from early embryogenesis were shown. Same legend as in Fig. 1D except the ATAC-seq from 60hr embryos is shown in green. **E)** Simulation of END-seq pattern using telomere addition sites. Shown are observed *Ascaris* END-seq data (top) compared to simulated END-seq profiles (middle) using the END-seq profiles from *O. tipulae*, a nematode with homogeneous genetic background and homogeneous DSBs. The simulation used the position and frequency of *Ascaris* telomere addition sites (bottom) (see Methods). Note the similarity between the observed and simulated END-seq profiles on the retained ends. **F** and **G**) Longer resection occurs in *Ascaris* compared to *O. tipulae*. **F)** Distance between the median resected end from the retained and eliminated sides. The median values for all *Ascaris* CBRs were plotted. All development times (54-98hr) are statistically significant compared to 50hr (significance is only shown between 50hr and 54hr). **G)** Distance between the median resected end from the retained and eliminated sides of the SFE in *O. tipulae*. Wilcoxon statistic tests were used with ** p < 0.01, **** p < 1e-4.

### Comparative analysis of CBRs between *Ascaris* and *Parascaris*

To examine sequence conservation among CBRs within *Ascaris* or *Parascaris*, the CBR sequences were compared against each other using *blastn* (-evalue 0.01) (38). Two CBRs were considered to have high sequence similarity if over 50% of the query CBR in length had a BLASTn hit to the subject CBR. The sequence conservation was also assessed between the CBRs from *Ascaris* and *Parascaris*. However, due to diverged sequence between the two species, the comparison was carried out at the translated amino acids level using *tblastx* (-evalue 0.01) (38). Two CBRs were considered to have high sequence similarity if over 50% of the query CBR had a tBLASTx hit to the subject CBR in the other organism. Random genomic regions (1000, 8 kb regions generated by BEDTools *random* -n 1000 -l 8000 -seed 123) were used to assess the overall sequence conservation between *Ascaris* and *Parascaris*, using tBLASTx of random region against the other species’ genome.

### Genome sequencing and analysis on X-ray irradiated embryos

The same method described for END-seq *de novo* telomere analysis was used to analyze the telomere addition events in the control vs. the irradiated embryos. END-seq reads with two consecutive telomeric repeat units were mapped to the genome. BEDtools *map* (-c 4 -o sum -null 0) was used to assess the number of reads in the CBRs, eliminated DNA region, and retained DNA region. The number of reads was normalized to the size of these genomic regions in kilobases (telomere reads/kb).

## RESULTS

### PDE-induced DSBs occur before mitosis in *Ascaris*

Previous genomic analyses in *Ascaris* somatic cells (comma stage embryos [7-day], post PDE) identified sites where new telomeres are added. These sites reside within a 3-6 kb genomic region known as a chromosomal breakage region (CBR) (24). However, the telomere addition sites do not necessarily correspond to the sites where the DSBs occurred during PDE, since following DSB induction, the DNA ends could be trimmed (removal of nucleotides at both strands) prior to the addition of telomeres. To identify the sites of the DSBs and their timing during the cell cycle of PDE, we used END-seq (31,32), a method based on the direct ligation of a sequencing adapter to the ends of DSBs after removing single-strand nucleotides, to capture DSBs and their end processing (resection). We first demonstrated that END-seq can identify exogenously introduced DSBs at single nucleotide resolution using a restriction enzyme (FseI) on *Ascaris* embryos (Fig. 1A and 1B). We observed that the END-seq reads are highly enriched at the junction of both the retained and eliminated DNA, indicating that END-seq can capture the endogenous DSBs associated with PDE (Fig. 1A).

In *Ascaris*, five independent PDE events occur in pre-somatic cells during the 4-16 cell stages, with four of them at the 4- or 8-cell stage (Fig. 1C). In the 4-cell embryo, two cells (ABa and ABp) simultaneously undergo PDE, followed by PDE in the EMS cell (39). Notably, *Ascaris* early embryos have a long cell cycle of ∼15 hours (40), compared to ∼20-30 minutes in the free-living nematodes *C. elegans* and *O. tipulae*. The long cell cycle allowed us to identify and examine DSBs and their resection at 11 time points between 50 to 98 hours of embryo development. This time frame covers discrete phases of the cell cycle during the 4-8 cell stages (40) (Fig. 1C). Our END-seq data indicated that DSBs for PDE were not detected during the S phase (50hr) of the 4-cell embryos. However, a small but significant amount of END-seq reads appear in the CBRs at 54hr (G2 phase), and the END-seq signals increase through 80hr (Fig. 1D-F). The initial detection of END-seq signals suggests the DSBs occur during the G2 phase of the cell cycle, prior to chromosome condensation and mitosis. The timing is different from a prevailing view that DSBs may occur during mitosis (41) and has important implications for the molecular mechanisms of DSBs (see Discussion).

### *Ascaris* DSBs are heterogeneous and undergo resection

Our END-seq data were derived from a heterogeneous population of millions of *Ascaris* embryos obtained from wild isolates. We reasoned if the DSBs were homogeneous in this population and occurred at a single location within a CBR, the END-seq would result in no overlapping reads from the two strands (see Fig. 1B). However, we observed all 72 CBRs have overlapping reads from the two strands (Fig. 1D-E and Fig. S1), suggesting DSBs occur heterogeneously within these overlapping regions in the sampled population. The overlapping regions of the END-seq signal coincide with the telomere addition sites within the CBRs, indicating telomere healing may occur at the site of the DSB without DNA trimming (see below for additional evidence). Overall, our END-seq data from a wild population of *Ascaris* embryos indicates that DSBs are heterogeneous, confined within the CBRs, and are likely the sites of telomere addition.

Although the DSB sites are heterogeneous in the population, the END-seq reads accumulate across extended regions of the CBRs on both strands and there is a large offset between the majority of retained and eliminated END-seq reads, as indicated in the distance between peaks of retained and eliminated ends (Fig. 1D-E), suggesting extensive bi-directional end resection from 5’ to 3’, leaving an extended 3’-overhang. Quantification of the END-seq reads revealed greater resection of eliminated ends of DSBs compared to retained ends of DSBs (Fig. 1F and Table S1). The resection bias is observed from 54hr to 80hr of embryo development. Because each DSB produces a retained and an eliminated end, this indicates the eliminated DNA ends are accessible longer than the retained ends for resection (and thus END-seq detection). This is consistent with the overall longer tails of END-seq reads observed on the eliminated side of the DSB, indicating continued resection (Fig. 1D-E and Fig. S1). After 80hr, we observe an overall decrease of END-seq signal (Fig. 1F), likely due to the completion of the first three PDE events and the one new PDE event at the 8-cell stage (see Fig. 1C). More but not significant amounts of reads were observed on the eliminated side (Fig. 1F). This more balanced number of reads between the eliminated and retained ends could be due to 1) the overall diminishing of END-seq signal in the eliminated regions from the previous three PDE events and 2) the early time point of the new PDE event where bias in resection has not accumulated. In sum, our END-seq experiments show that the DSBs occur during the G2 phase of the cell cycle; they occur heterogeneously within the CBRs; and the DSBs undergo bi-directional resection.

### *Ascaris* DSB end resection generates 3’ overhangs with site-specific patterns

To further characterize the ends of DSBs associated with PDE, we sought to determine the percentage of *Ascaris* END-seq reads that are blunt vs. have an overhang. Given that the standard END-seq procedure can capture DSBs with both blunt ends and 3’ overhangs (31,32), we modified END-seq to capture 1) only blunt ends (direct capture) or 2) blunt ends, 3’-, and 5’-overhangs (all-END) (Fig. 2A). Our direct capture method identified fewer reads in the CBRs compared to the all-END method (Fig. 2B-C). The blunt-end reads identified through the direct capture method are likely derived from the initial DSBs that have not undergone end resection. They are largely confined within the CBR, further supporting that the sites of DSBs are confined within the CBR and are the sites of telomere addition. Quantification of the END-seq reads from all CBRs suggests that 79% correspond to resected DSBs with a 3’-overhang (Fig. 2C). In addition, analysis of the resection profiles between individual CBRs reveals notable differences in the frequency of END-seq reads and resection endpoints (Fig. 2D and Fig. S1). However, within a specific CBR, the resection profile was highly consistent across all developmental stages (Fig. 2D and Fig. S1). The variations in resection profiles among CBRs could be due to the local sequence, nucleosome organization, and chromatin structure (see ATAC-seq data in Fig. 2D and Fig. S1) that may influence the resection process and endpoints, as illustrated in recent studies (42,43). Overall, our analysis revealed that most DSBs are resected to generate long 3’ overhangs, and the resection profiles are site-specific.

### *De novo* telomere addition occurs at the DSB site

Our previous END-seq analyses on *O. tipulae* PDE revealed that DSBs occur at the center of a 30 bp, degenerated palindromic sequence (SFE) and that telomeres are added at the sites of DSBs (20). We wondered if telomere addition sites are similarly close to the DSBs in *Ascaris*. However, the heterogeneity of telomere addition sites from the wild population of *Ascaris* embryos makes it difficult to directly assess a single breakage event. We thus used a computer simulation to indirectly evaluate the likelihood of telomere addition sites corresponding with DSB sites. In this simulation, we applied the average *O. tipulae* END-seq resection profile to each observed telomere addition site in *Ascaris* (considering both the position of the telomere site and its frequency; see Methods). Interestingly, the simulated END-seq profiles of the retained ends match consistently with the observed *Ascaris* END-seq data, suggesting telomere addition sites likely correspond with the DSB sites (Fig. 2E and Fig. S2). Together, several lines of evidence suggest that *de novo* telomere addition occurs at the DSB site in *Ascaris*, including 1) the overlapping regions of END-seq signal where DSBs presumably occur, coinciding with the telomere addition sites (Fig. 1D-E), 2) the blunt-end reads are confined within the CBR, where telomeres are added (Fig. 2B), and 3) the simulation shows consistent END-seq profile between *O. tipulae* and *Ascaris* on the retained side of DSBs (Fig. 2E).

**Figure 3.**
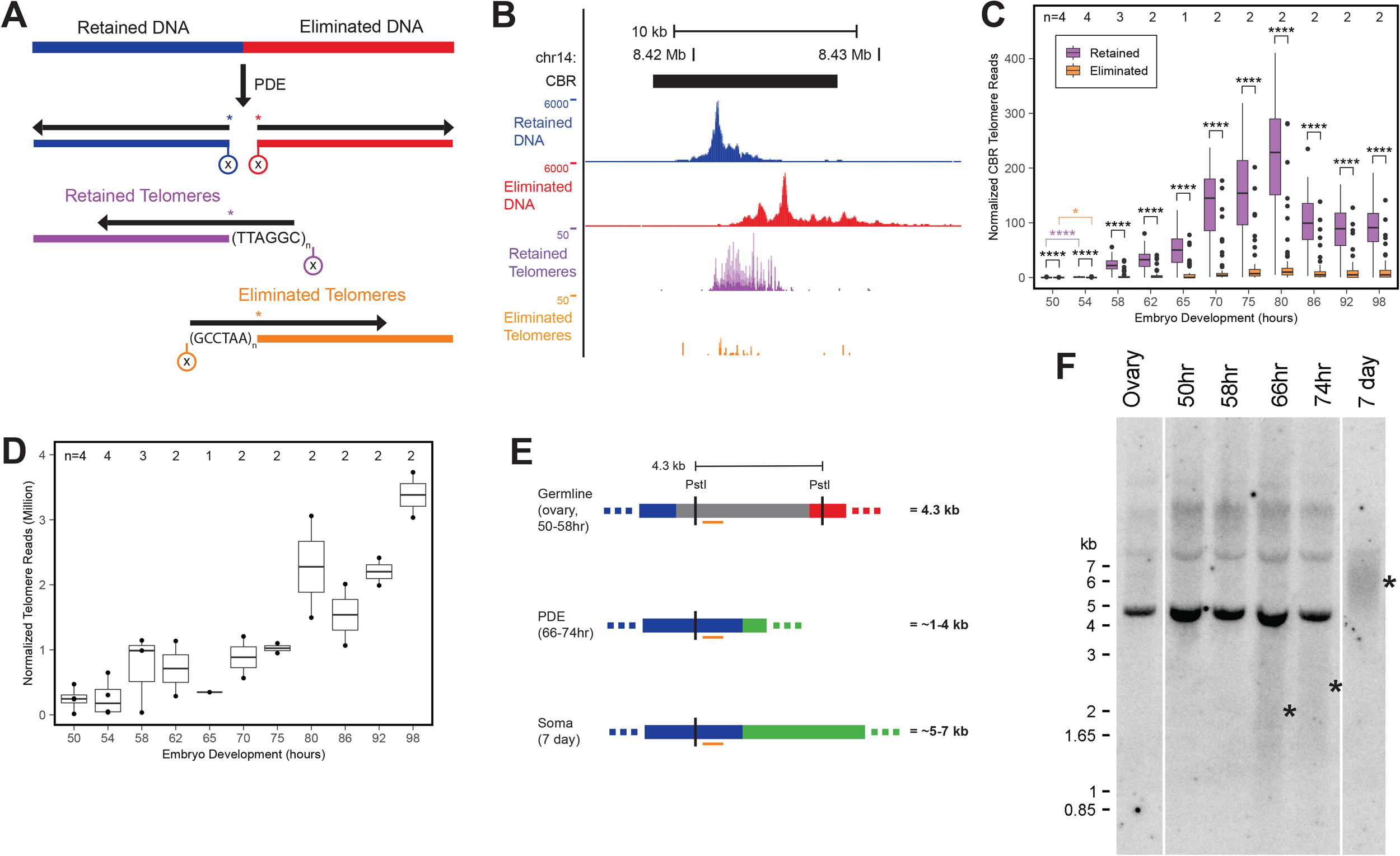
Telomere addition in *Ascaris* favors the retained ends of DSBs. **A)** A schematic showing the sequence ends with and without *de novo* telomeres captured by END-seq. Blunt and resected DSB ends were trimmed and captured with END-seq (circle with X, blue/red for ends without telomeres, and purple/orange for ends with new telomeres, not drawn to scale). The horizontal black arrows indicate END-seq reads pointing from 5’ to 3’. For ends without telomeres, the 5’-ends of the reads (first nucleotide captured, asterisk) were used for data analysis. New telomeric sequences (TTAGGC/GCCTAA)n with their length shorter than the length of sequencing read (150 bp) are indicated. The unique (non-telomeric) region of the reads was mapped to the genome, with the first non-telomeric base (5’) designated as the telomere addition site (asterisk). **B)** Majority of the telomere addition occurs at the retained ends. A genome browser view of the two types of END-seq reads (split by strand into four tracks, see Fig. 3A) captured by END-seq at a CBR. **C)** Biased telomere addition is consistent across all CBRs and developmental stages. Average END-seq telomere signal in each CBR across development. At each time point, there are significantly more telomere reads from the retained side of the DSB. All time points also have significantly more retained and eliminated reads than 50hr (significance only shown between 50hr and 54hr). **D)** The number of telomere-only END-seq reads plotted across development. For Fig. 3C-D, the number of biological replicates is indicated at the top of the graphs. Wilcoxon test * p < 0.05, **** p < 1e-4. **E)**. A schematic of the Southern blotting. On the left is a CBR from chromosome 6 (CBR_m6b), with the restriction sites and region for the probe. Blue = retained DNA, gray = CBR, red = eliminated DNA, green = new telomere, vertical lines = PstI sites, and orange horizontal bar = 700 bp probe region. On the right is the predicted size of the DNA in the sampled tissues or developmental stages. **F)**. Southern blot showing the intact germline DNA (4.3 kb) and the various sizes of DNA in different embryonic stages. The * indicates the estimated average size of somatic DNA hybridized to the probe. Note the gradual increase of the somatic DNA size with development.

However, for the eliminated ends of the DSBs, the simulation does not match the observed END-seq profiles (Fig. 2E and Fig. S2). Instead, the broken ends underwent much longer resection compared to the simulated profiles. This reflects differences in the resection of eliminated ends between *Ascaris* and *O. tipulae*. To further compare resection between the two nematodes, we analyzed the distance between the median retained and eliminated ends. We found *Ascaris* has a much longer median distance (majority 1-3 kb) compared to *O. tipulae* (majority < 0.5 kb, see Fig. 2F-G). Since the resection profiles on the retained sides appear consistent between *Ascaris* and *O. tipulae* (Fig. 2E and Fig. S2), this indicates the length of resection on the retained sides is largely the same between these nematodes. Thus, the observed difference in distance may be caused by the extended resection at *Ascaris* eliminated ends (Fig. 1D-F). In addition, the heterogeneous nature of DSBs within the *Ascaris* population contributes to the longer resection distance. Overall, the consistency of resection profiles on the retained ends suggests the mechanism of end resection is likely conserved between *Ascaris* and *O. tipulae*, while longer resection at the eliminated ends indicates an extended processing of DSB ends of the eliminated DNA in *Ascaris* (see below).

### Telomeres are preferentially added to retained DNA ends

In a previous study, the telomere addition sites were defined using genome sequencing on comma-stage (7-day) embryos long after the PDE events (24). While their positions in the genome were determined for the retained ends, little is known about telomere addition at the eliminated sides since the sequences are absent in the comma-stage embryos. In addition, the timing and speed of telomere addition were also not known during PDE. Using PCR amplification, Jentsch et al. showed that telomeres can be added to both retained and eliminated ends in *Ascaris*, suggesting telomere addition may be a non-specific process (44). More recently, in *O. tipulae*, we showed telomeres are added to both broken ends in an unbiased manner (20), consistent with a non-specific telomere healing model. Here, we assessed *Ascaris de novo* telomere addition at both the retained and eliminated ends by extracting and analyzing telomere-containing reads from our END-seq data (Fig. 3A). Surprisingly, our data showed telomeres are primarily (overall 89% of their reads) added to the retained ends of DSBs (Fig. 3B and Table S1). Our data is consistent with the previous work in *Ascaris* (44) since 11% of telomere addition on the eliminated ends would still allow its detection by PCR. However, this result contrasts with the unbiased telomere addition observed in *O. tipulae* (20), suggesting a molecular difference between these nematodes (see discussion).

We further assessed *de novo* telomere addition across development to determine its timing and extension through PDE. Overall, the ratio of telomeric to non-telomeric END-seq within the CBRs suggests most (97%) of the DSBs are not readily healed with telomeres, likely an indication of active processing (resection). For the 3% telomeric reads, we found a striking similarity of the profile of change through development (Fig. 3C) compared to the non-telomeric reads (see Fig. 1F), suggesting a small portion of telomere addition can happen with little or no lag time after formation of DSBs. Importantly, END-seq can only map added telomeres to a unique site when the telomere length is shorter than the sequencing reads (150 bp in our Illumina sequencing). To account for all telomeres, we quantified the number of END-seq reads that contained two or more consecutive telomeric repeat units (Fig. 3D and see Methods). These telomeric reads rise steadily from 50hr to 75hr, likely due to the growth of new telomere ends that are greater than 150 bp. Interestingly, we found a dramatic rise in these telomeric reads during the 75hr to 98hr time points. The number of telomeric reads is much higher than expected if we only consider the increase caused by karyotype changes that occur during PDE (increase from 24 germline chromosomes to 36 somatic chromosomes). We interpret this increase in telomeric reads as the result of the fragmentation of old germline telomeric sequences during their degradation, leaving numerous small telomere fragments captured by END-seq. In agreement with this, analysis of these telomeric reads in the 75-98 hours indicates that the majority (59%) of them are telomere-only reads. Overall, these data show the timing of germline telomere breakdown and somatic telomere synthesis during PDE.

To further corroborate the timing of telomere addition, we performed Southern blotting using a probe targeted to a single CBR (Fig. 3E). This allows us to determine changes to the DNA at the CBR in the germline, during early embryos through PDE stages, and in somatic cells. The result confirms the CBR is intact (4.3 kb DNA band) in the germline (ovary), while almost all DNA at this CBR was broken in the somatic cells (7-day embryos, about 500 somatic cells with two primordial germ cells) (Fig. 3F). The smear observed in the somatic cells indicates a heterogenous length, likely caused by the different DSB sites and variations in the length of newly added telomeres (Fig. 3E). During PDE (66hr), we observed in addition to the 4.3 kb germline DNA, a smear of DNA with peak density at 1-4 kb, suggesting a broken CBR with heterogenous break sites (and telomere lengths) in the population of embryos (Fig. 3F). Although it is difficult to quantify the amount and the exact length of the DNA smear, we observed a clear shift of the smear toward larger DNA in 74hr, reflecting an increase in telomere length (Fig. 3F). In sum, our *de novo* telomere addition analysis provides insights into the timing, selection, and dynamics of telomere addition during *Ascaris* PDE.

### Alternative break sites provide a fail-safe mechanism for PDE in *Ascaris*

Previous genomic studies revealed 72 *Ascaris* CBRs (canonical CBRs, or canCBRs) - defined by their genomic positions at the junction of retained and eliminated DNA, where new telomeric sequences are detected in somatic cells (19,24). These studies did not determine if DSBs and telomere addition also occur within the eliminated regions or how the eliminated DNA is degraded. Here, we identified 28 additional break regions in the eliminated DNA, hereafter called alternative CBRs (altCBRs, see Fig. 4A and 4B). These alternative CBRs appear to occur simultaneously with the 72 canonical CBRs based on a similar number of END-seq reads; undergoing bi-directional resection and are healed with *de novo* telomere addition at a low level, similar to eliminated ends of the CBRs (Fig. 3B). These alternative CBRs could serve as a fail-safe mechanism to ensure PDE occurs, as seen in *O. tipulae* (20). Since the assembled *Ascaris* genome is not telomere-to-telomere, sequences in the eliminated regions are incomplete and some regions contain highly repetitive elements (19). Therefore, we reason additional alternative CBRs may exist but were missed in our analysis. Interestingly, many of the alternative CBRs were found in internally eliminated sequences (19 of 28, 68%), which consist of only 42% of all eliminated DNA. These DNA sequences are between evolutionarily fused chromosomes (Fig. 4B) and may suggest a critical role of PDE in breaking the chromosomes to restore their pre-fused karyotypes (21). We further compared the conservation of sequence between all CBRs to determine their relationships and evolution. One large and two small groups of CBRs showed high sequence similarity (Fig. 4B and Table S2), suggesting these *Ascaris* CBRs have been recently duplicated, similar to the alternative break sites observed in *O. tipulae* (20). Interestingly, we observed a high number of canonical-to-canonical and alternative-to-alternative pairs but a low number of canonical-to-alternative pairs (Fig. 4C and Table S2), suggesting some constraints on the interchangeability of the canonical and alternative sites. Nevertheless, the presence of alternative CBRs as a potential fail-safe mechanism for PDE further suggests the biological importance of PDE in *Ascaris*.

**Figure 4.**
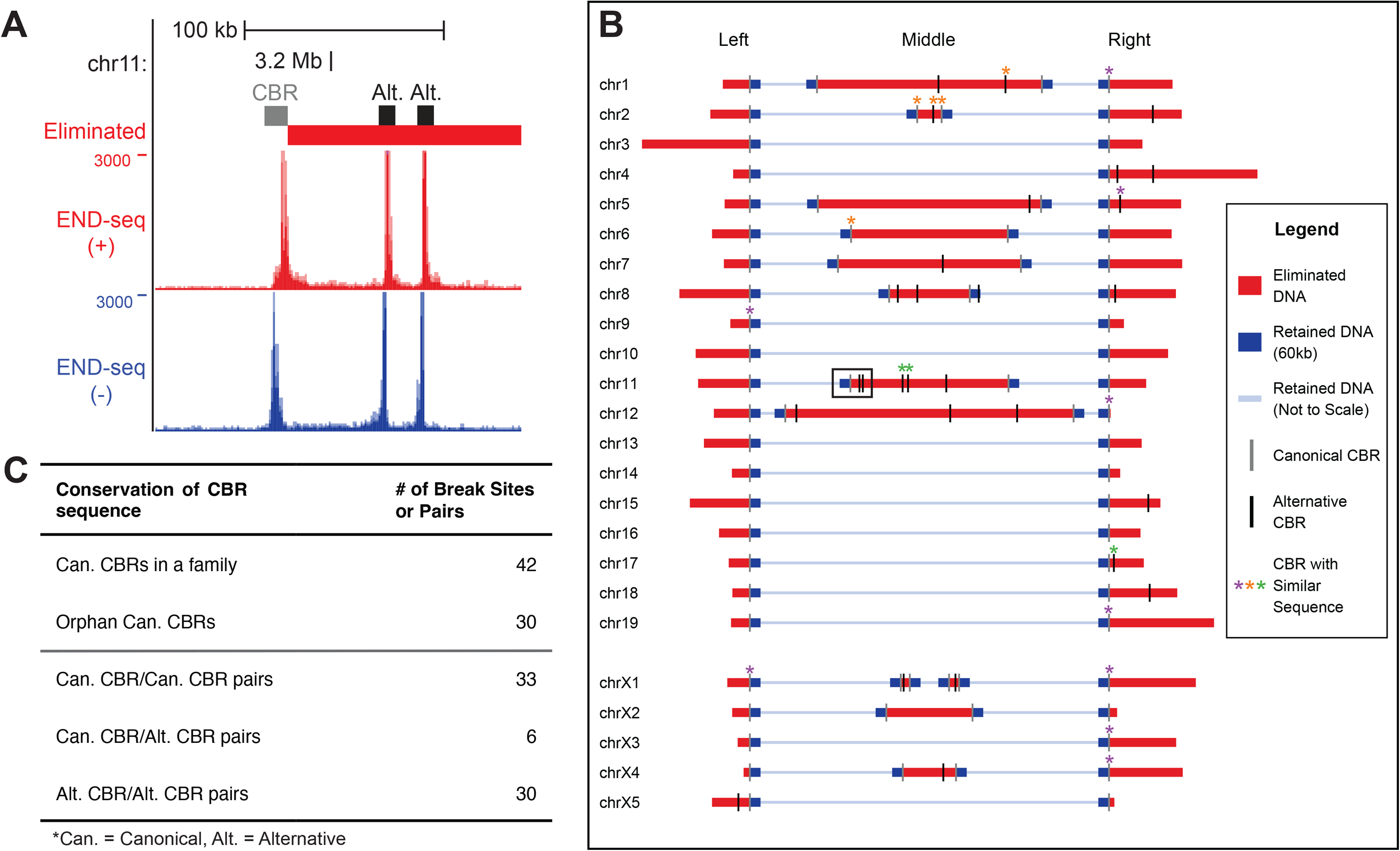
Alternative CBRs in *Ascaris* suggest a fail-safe mechanism for PDE. **A)** END-seq reveals alternative CBRs in the eliminated regions. A genome browser view of a canonical CBR and two alternative CBRs within the eliminated DNA. **B)** Distribution of canonical CBRs and alternative CBRs in the *Ascaris* genome. A schematic showing the position of all *Ascaris* CBRs. The region shown in **A** is indicated with a black box. To emphasize the eliminated DNA, most sequences of a chromosome are represented by a thin, pale blue line not plotted to scale. Eliminated DNA (red) and 60 kb flanking retained DNA (blue) are plotted as thick lines and drawn to scale. Asterisks mark clusters of CBRs (3 or more CBRs) that have >50% nucleotide sequence identity. **C)** A summary table of nucleotide sequence identity among CBRs.

### Telomere addition is specifically linked to PDE-induced DSBs

Our data indicates that all retained PDE-induced DSB ends are healed with telomere addition (Fig. 1 and S1). We wondered if this healing is specifically linked to PDE or occurs universally in all DSBs generated during the PDE stages. We irradiated *Ascaris* early embryos undergoing PDE with 100-200 Gy of X-ray irradiation to introduce exogenous DSBs, followed by END-seq and genome sequencing to evaluate telomere addition across the genome. Irradiated embryos showed a significant developmental delay after 24hr and 48hr post-irradiation (Fig. 5A and Table S3). While direct detection of exogenous DSBs in these *Ascaris* embryos was difficult, the observed developmental delays (Fig. 5) suggest an impact from the X-ray treatment (45). However, the embryos were able to recover, continue to develop, and progress through cell cycles as we observed an increased number of cells between irradiation and 48hr of recovery. If X-ray-introduced DSBs are also healed with telomere addition, we would expect to see an increase in telomere-containing reads across the genome. Our END-seq data on control embryos showed that telomere addition occurs mostly within the CBR regions, and very few telomere reads were found in the eliminated regions and almost none in retained regions (Fig. 5B). This END-seq result is consistent with the genome sequencing data and suggests our method can capture telomere addition across the genome. However, our END-seq on irradiated embryos revealed no significant increase in telomere addition within the retained or eliminated genome regions (Fig. 5B). Future experiments on the impact of the X-ray, including the sites and amounts of DSB and how they may be repaired are needed. Nevertheless, our data suggests that PDE-induced DSBs are specifically marked for telomere addition.

**Figure 5.**
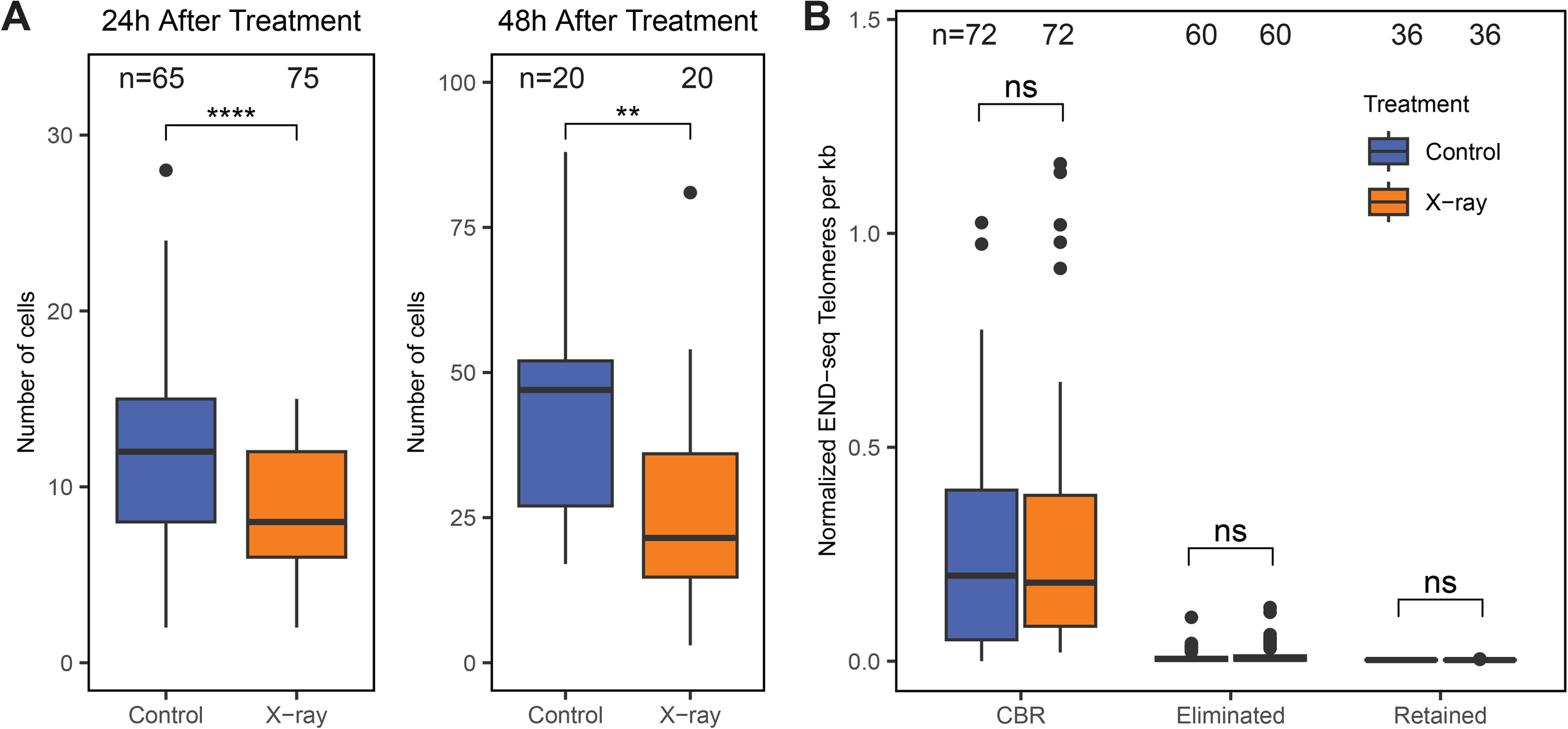
Telomere addition is specific to PDE-induced DSBs. **A)** X-ray-treated embryos (4-cell, 65hr) show delays in their development. Number of cells in each embryo compared between irradiated (100 Gy X-ray) and control cells. Embryos were allowed to recover for 24 or 48 hours before cells were counted. **B)** Telomere addition was not detected in non-CBR genomic regions in X-ray-treated embryos (4-6-cell, 70hr). Number of END-seq telomeres/kb found in each genome region from control and irradiated cells. T-test ** p < 0.01, **** p < 1e-4.

### End resection and telomere addition are conserved in the horse parasite *Parascaris*

A closely related parasitic nematode from the horse, *Parascaris univalens*, also undergoes PDE (39). We performed END-seq in *Parascaris* early embryos and compared it to *Ascaris*. Overall, our *Parascaris* data showed a close resemblance to observations in *Ascaris*. *Parascaris* DSBs undergo extensive bi-directional resection, with the eliminated ends undergoing longer resection than retained ends, as well as a biased telomere addition towards retained ends (Fig. 6A). Furthermore, 27 alternative CBRs (6 in unplaced contigs) were identified. To investigate the divergence of the CBR sequences and their potential rearrangements within the chromosomes, we compared the CBR sequence and their positions in *Ascaris* and *Parascaris* genomes. Interestingly, only about half (34/72, 47%) of the canonical CBRs have sequence similarity among CBRs between the two species, and about 68% of the CBRs have a match across the entire genome of the other species (Fig. 6B-C and Table S2), while in comparison, about 93% of the randomly selected genomic regions can be matched between *Ascaris* and *Parascaris* across the genome (see methods and Table S2). This suggests the CBR sequences are fast-evolving regions of the genome. Interestingly, one CBR in *Parascaris* appears to have diverged into seven CBRs in *Ascaris* since the split of these species. Notably, the alternative CBRs appear to be the least conserved CBRs between the species, supporting a model that the eliminated DNA is more flexible and may be undergoing more rapid evolution (1,46,47). In sum, while the closely related *Ascaris* and *Parascaris* share many PDE features, there are notable variations in the sequences of the CBRs and their positions in the chromosomes, suggesting flexibility in the genomic location and the amount of sequence eliminated in nematode PDE.

**Figure 6.**
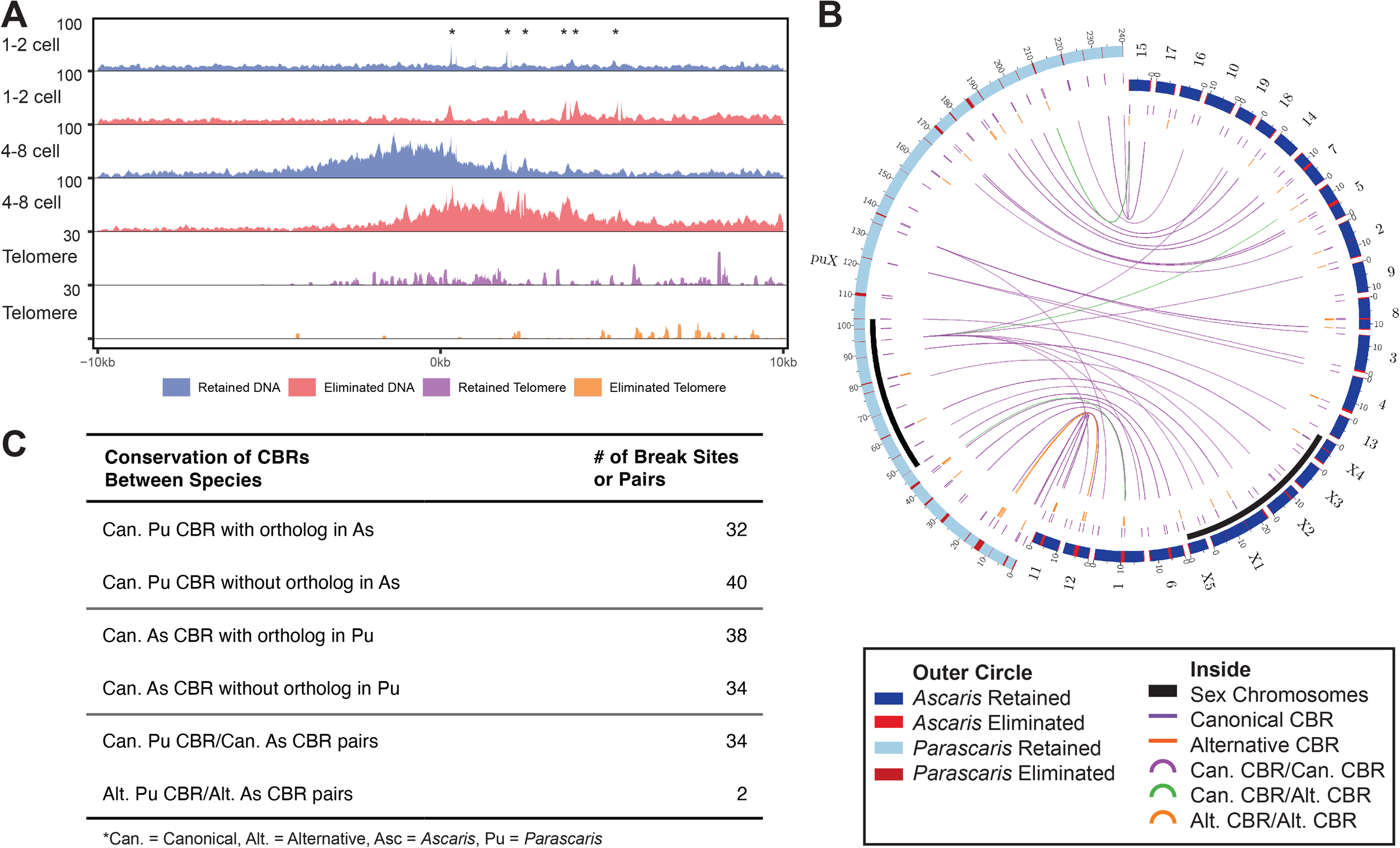
End resection and telomere addition in *Parascaris*. **A)** End resection profiles and telomere addition are similar in *Ascaris* and *Parascaris*. The average END-seq profile (100 bp bins, 10 bp sliding window, 20 kb). All 72 *Parascaris* CBRs were aligned by the median somatic telomere addition site. Asterisks mark background END-seq signal from repetitive sequences. **B)** Conservation of CBRs between *Ascaris* and *Parascaris*. A circos plot showing sequence similarity between *Ascaris* and *Parascaris* break sites. The outer circle is colored by eliminated (red) and retained (blue) DNA. Inside, for the next two tracks, purple lines indicate canonical CBRs and orange lines indicate alternative sites. Links connect CBRs with >50% sequence identity (defined by tBLASTx), with purple links connecting canonical CBRs, orange links connecting alternative sites, and green links connecting canonical CBRs and alternative CBRs. **C)** A summary table of sequence identity among CBRs with tBLASTx.

## DISCUSSION

Most unscheduled DNA double-strand breaks (DSBs) are harmful because failure to repair these DSBs compromises the integrity of the genome. However, controlled formation of DSBs is integral in some biological processes, such as the V(D)J recombination in immune cells (48,49) and homologous recombination during meiosis (50). Controlled DSB formation is also necessary in some organisms undergoing programmed DNA elimination (PDE), where chromosomes are fragmented, and DNA sequences are lost (1–4). In nematode PDE, little is known about what causes the DSBs and how the broken ends are processed. Here, we used END-seq on staged embryos and carried out in-depth analyses of DSBs during *Ascaris* PDE. We propose a model (Fig. 7) to describe the *Ascaris* DSBs for PDE, their end processing, and telomere addition and how these processes may differ from PDE in the free-living nematode *O. tipulae*.

**Figure 7.**
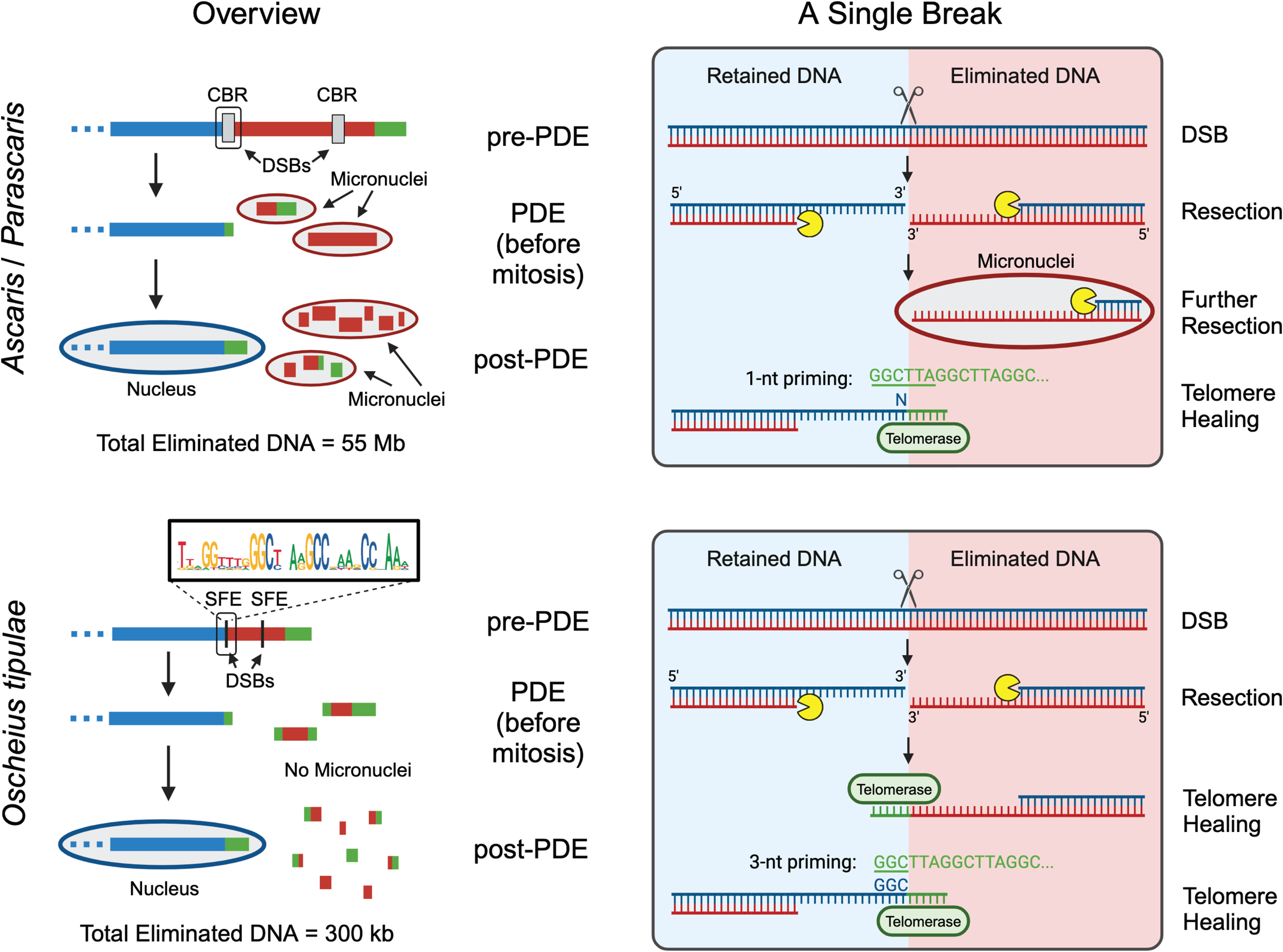
DSB, end resection, and telomere healing during nematode PDE. A comparison of DSB, end resection, and telomere healing between *Ascaris* and *O. tipulae*. Left (Overview): A model of PDE at the chromosomal level, not drawn to scale. Blue, red, and green rectangles represent retained, eliminated, and telomeric DNA, respectively. Ovals with a blue outline are nuclei and ovals with a red outline are micronuclei. For *Ascaris* and *Parascaris* (upper left), DSBs occur within a 3-6 kb CBR (grey box) and remove a total of a total of 55 Mb (*Ascaris*, 18%) and 2.2 Gb (*Parascaris*, 90%) of DNA from the genome. Some eliminated regions also contain alternative CBRs. Retained DNA is healed with *de novo* telomere addition while eliminated DNA is not. The retained DNA is selectively segregated to the nuclei. In contrast, eliminated DNA is encapsulated in micronuclei where they are further resected and eventually degraded. For *O. tipulae* (Bottom left), DSBs form at the center of a 30 bp SFE motif (boxed consensus sequence; vertical black line marks SFE position) and remove a total of 350 kb (0.6%) of the genome. Some eliminated regions contain alternative SFEs which act as a fail-safe mechanism. After DNA break formation, both retained and eliminated sequences are healed with *de novo* telomere addition. Right (A Single Break): A model of PDE at one break site, not drawn to scale. The scissors represent a presumptive nuclease that generates a DSB, and the Pac-man represents exonucleases involved in end resection. Micronuclei are shown as red ovals. New telomeres are represented in green). For *Ascaris* and *Parascaris* (upper right), a DSB is generated at a single spot within the CBR and undergoes bi-directional resection, generating a long 3’-overhang. The retained end of the break (left, blue region) is healed with *de novo* telomere addition, while the eliminated end of the break (right, red region) is encapsulated in a micronucleus and continues to undergo resection. Telomeres are added directly to the site of the retained DNA break. *Ascaris* uses 1-nt priming where any nucleotide can prime telomere addition. For *O. tipulae* (bottom right), a DSB is generated within the SFE and undergoes bi-directional end resection, generating a long 3’-overhang. Both retained and eliminated ends are healed with *de novo* telomere addition. Telomeres are added directly to both sides of the DNA break, likely using the conserved GGC for telomerase priming. Figure created with BioRender.com.

### Timing of DSBs

Our data revealed that DSBs occur during the G2 phase of the cell cycle (Fig. 1). This indicates the chromosomes are broken before the onset of mitosis. We previously showed that *Ascaris* chromosomes are holocentric, and before and during the mitosis for PDE division, the to-be-eliminated DNA is devoid of centromeres (51). Electron microscopy (EM) showed that during an *Ascaris* PDE mitosis, fragments of chromosomes that will be eliminated do not align at the metaphase plate and they lack kinetochores and microtubules (19), supporting our END-seq data that the chromosomes were already broken before metaphase. The DSBs likely occur while the chromosomes are decondensed, and the CBRs are accessible to machinery that may generate, process, and/or repair the DSBs. This is consistent with our ATAC-seq data that the chromatin at the CBRs is more open during and after PDE (24). Potential mechanisms that lead to the DNA breaks may include the formation of R-loops (52–54) and interactions of CBRs at the 3D genome level (55–57). These mechanisms are not dependent on the presence of a sequence motif, in agreement with the heterogeneous and sequence-independent DSBs and telomere addition sites observed in *Ascaris* (19,24).

### DSBs, end resection, and telomere addition

The heterogeneous telomere addition sites observed within a CBR in the *Ascaris* embryo population could each be derived from a single DSB site that is trimmed to variable lengths before the addition of new telomeres. However, several lines of evidence indicate trimming is unlikely. First, trimming from a single site would leave a gap between the two broken ends, resulting in a lack of END-seq signal in the gap region. We did not identify any gap in the END-seq signal within the CBR. Second, we reason the sites where the blunt-end reads have not undergone resection are likely the sites where the DSBs originated. These sites coincide with the telomere addition sites (Fig. 2B), suggesting no trimming is needed before telomere healing. In addition, we observed consistency between the retained side of our END-seq data and the simulation profile (Fig. 2E) from *O. tipulae* in which the sites of *de novo* telomere addition are the sites of DSBs (20). Together, these data suggest that the DSB is not trimmed but undergoes resection from 5’ to 3’, leaving an extended 3’-overhang where the new telomere is primed and added at the site of the DSB (Fig. 7). We believe this 3’-overhang structure provides a readily accessible substrate that facilitates *de novo* telomere healing via telomerase (58,59), and the resected nucleotides will be filled with lagging-strand synthesis and telomere C-strand fill-in likely through CST–polymerase α-primase (60,61). In yeast and human, extensive 5’ to 3’ resection activates Mec1/ATR-dependent signaling which blocks telomerase from converting DSBs into telomeres (62,63). It is plausible that in nematodes, the end resection of PDE DSBs may be repressed when the telomere maintenance machinery (telomerase and CST–Polα/Primase) acts on the DNA substrate. The resection profiles appear to reflect the positions and frequencies of telomere addition (Fig. 3), and each profile is likely shaped by the local sequence and chromatin features (42,43).

The broken chromosome fragments would need to be protected until the telomere healing occurs. Our END-seq data showed only a small percentage (3%) of the captured ends within CBRs have telomeres, suggesting most of the DSBs are undergoing end resection during the initial stage of PDE. We speculate the resection machinery may be tightly linked to the DSB break generation. This association could be achieved through specific foci or condensates organized within the nuclei where enzymes for the breaks, end processing, and repair are enriched. This is consistent with our preliminary observation that CBRs are interacting with each other at the 3D genome level during the time of PDE (Simmons and Wang, personal communication). We reason the resected ends would prevent the NHEJ DNA repair pathway from acting on the ends since resection of DSBs is thought to inhibit NHEJ (64). In contrast, the long resected end would be suitable for homologous recombination (HR) mediated repair (65). However, all our data show no sign of recombination or genome rearrangement during PDE, illustrating the broken DNA ends are consistently healed by telomere addition (19,24,25). This suggests either the HR pathway is not active, unavailable, or out-competed by the telomere maintenance pathway (Fig. 7). Future studies are warranted to elucidate mechanisms of the choice of DNA repair pathways for the DSBs during PDE.

### Biased telomere addition and micronuclei

To our surprise, even though end resection happens bi-directionally to both the retained and eliminated ends, 89% of the telomere addition events occur only to the retained ends of the DSBs (Fig. 3). This may indicate the eliminated ends are not available or accessible to the telomerase. Our previous EM data showed that the DNA fragments to be eliminated were engulfed into micronuclei (19). The rapid sequestration of eliminated fragments into micronuclei and the time required for telomere addition to occur may restrict the addition of telomeres to the eliminated DNA, leading to biased telomere addition only at retained ends. The micronuclei may not contain telomerase and thus telomeres are not added. However, end resection machinery appears present in the micronuclei as extended resection occurs on the eliminated sides of the DNA (Fig. 1D-E). A few telomere addition events to the eliminated sides likely happened before the DNA fragments were engulfed into the micronuclei, allowing them to be detected by PCR in a previous study (44) and END-seq in this work.

In contrast, we observed in *O. tipulae* that both the retained and eliminated sides have the same amount of telomere addition (20). The differences in the telomere addition to the eliminated ends may be due to the sequestration of the DNA into micronuclei in *Ascaris* and the loss of only small amounts of DNA in *O. tipulae* (Fig. 7). *O. tipulae* has a 60-Mb genome and eliminates only ∼0.6% (350 kb) of the DNA (20,66), while *Ascaris* has a genome of 308 Mb, and it removes 18% (55 Mb) of its genomic sequences (19). This to-be-eliminated DNA will be in the cytoplasm after the completion of mitosis. Cytoplasmic DNA can trigger a variety of cellular responses that can be deleterious to the cells (67–69). Thus, it may be critical to sequester, mask, or rapidly degrade the cytoplasmic DNA. The eliminated sequences in *Ascaris* persist for 2-3 cell cycles (∼50-60 hours) after PDE mitosis and are readily visible using DAPI staining (2,19,70). In contrast, the DNA in *O. tipulae* is not detectable using DAPI or Hoechst staining (20) and only takes 1-2 hours to degrade. Given the small amount of eliminated DNA and its short existence, *O. tipulae* may not need to sequester the eliminated DNA into micronuclei to prevent adverse effects (Fig. 7). Consistent with this model, *Parascaris* eliminates a large amount of DNA (2.2 Gb, 90% of the germline genome) and our END-seq data showed a biased telomere addition (Fig. 6), suggesting the eliminated DNA may also go into micronuclei. Further studies in additional nematodes with PDE may reveal the relationship between cell cycle length, the amount of eliminated DNA, the time of its degradation, its association with the formation of micronuclei, and its impact on telomere addition to the eliminated DNA.

### Comparison of DSBs and telomere addition among PDE species

The DNA substrate, telomeric sequence (TTAGGC), and mechanism of telomere addition appear the same in *Ascaris* and *O. tipulae* (Fig. 7). Both nematodes use extensive end resection to generate 3’-overhangs, and new telomeres are added to the DSB sites without trimming. The major difference in PDE between these nematodes, however, lies in the identification of DSB sites. In *O. tipulae*, a conserved motif (SFE) is required for the break (20), while in *Ascaris*, the DSBs are not associated with a specific sequence and can occur at any position within the CBR. This difference suggests divergent mechanisms for the recognition of the DSB sites and/or the generation of the breaks. In *O. tipulae*, it is plausible a DNA binding protein(s) may recognize the palindromic SFE motif, while in *Ascaris*, mechanisms independent of the sequences would be required to identify the CBRs.

This difference between the motif-based and the sequence-independent mechanisms may be associated with the variations in the sequence requirement in these nematodes for telomere addition (Fig. 7). In *O. tipulae*, the GGC sequences flanking the break are conserved across all SFE sites (20). This GGC matches the telomeric sequence TTA<u>GGC</u> and appears to be used for priming during telomere synthesis. It is plausible that *O. tipulae* requires this critical GGC for telomere healing, which could put evolutionary constraints on maintaining this sequence across all break sites. The constraint of this specific sequence may have co-evolved additional sequences surrounding the breaks, thus enhancing, and eventually fixing the use of the SFE motif. In contrast, sequence analysis showed that a single nucleotide of homology is sufficient for telomere addition *in vivo* in *Ascaris* (24). Since the *Ascaris* telomeric repeat, TTAGGC, contains all four bases, this allows the telomere to be added at any site within the CBRs; thus, there may be little or no evolutionary pressure to maintain any specific sequence for *Ascaris* telomere addition (Fig. 7). It would be interesting to carry out comparative analyses of PDE in more nematodes to further determine if the requirement for sequencing priming during telomere addition is connected to the usage of motif sequences for PDE breaks.

The motif-based and the sequence-independent DSBs in nematodes are reminiscent of PDE in ciliates, where in some species (*Tetrahymena* and *Euplotes*), specific motifs are used to generate the DSBs (71–73), while in others (*Paramecium*) the break sites appear to be sequence-independent (12). Interestingly, in *Tetrahymena*, the initial DSB ends are trimmed by a variable distance of 4-30 bp, leading to heterogeneity in the telomere addition sites despite using a motif-based mechanism (74). This differs from the telomere addition in *O. tipulae*, where telomeres are added directly to the break site (20). In contrast, in *Paramecium*, microheterogeneity (500-800 bp) and macroheterogeneity (several kbs) are observed for telomere addition (12), similar to the canonical CBRs and alternative CBRs observed in *Ascaris* and *Parascaris*. Telomerase is responsible for telomere addition in ciliates (12). We identified a single telomerase gene in *Ascaris* and *O. tipulae*, and its expression is elevated in both species during PDE (20,24,25,30), suggesting the telomerase is likely responsible for telomere addition during PDE. Overall, PDE in nematodes requires identification of the sites for DNA breaks, generation of the DSBs, and processing and repair of the broken DNA ends. However, the molecular features and the machinery involved in these processes appear to differ among diverse species, suggesting independent origins and evolution of these mechanisms.

## CONCLUSION

DSBs are harmful to the genome. They are mainly repaired by NHEJ and HR. An alternative repair pathway may be telomere addition. In ciliates and some nematodes that undergo PDE, telomere healing of DSBs is developmentally programmed, highly reproducible, and carefully regulated. Despite being known for over 130 years, little was known previously about the molecular details of DSBs and telomere healing processes in these parasitic nematodes. Our study provided insights into the timing of the DSBs, the dynamics of end resection, and the biases of telomere healing. The differential healing of retained vs. eliminated ends highlights a potential role of the micronuclei in confining the eliminated DNA. Our comparison also provides insights into the telomere priming and the sequence requirement for PDE. Future studies on nematode PDE may provide new insights into DSB generation, end processing, and telomere healing.

## DATA AVAILABILITY

The END-seq data is available at NCBI GEO with accession number GSE260958. The data is also available in a UCSC Genome Browser track data hubs that can be accessed with this link: https://genome.ucsc.edu/s/jianbinwang/ascaris_end_seq.

## SUPPLEMENTARY DATA

### List of supplementary Figures and Tables

**Figure S1.**
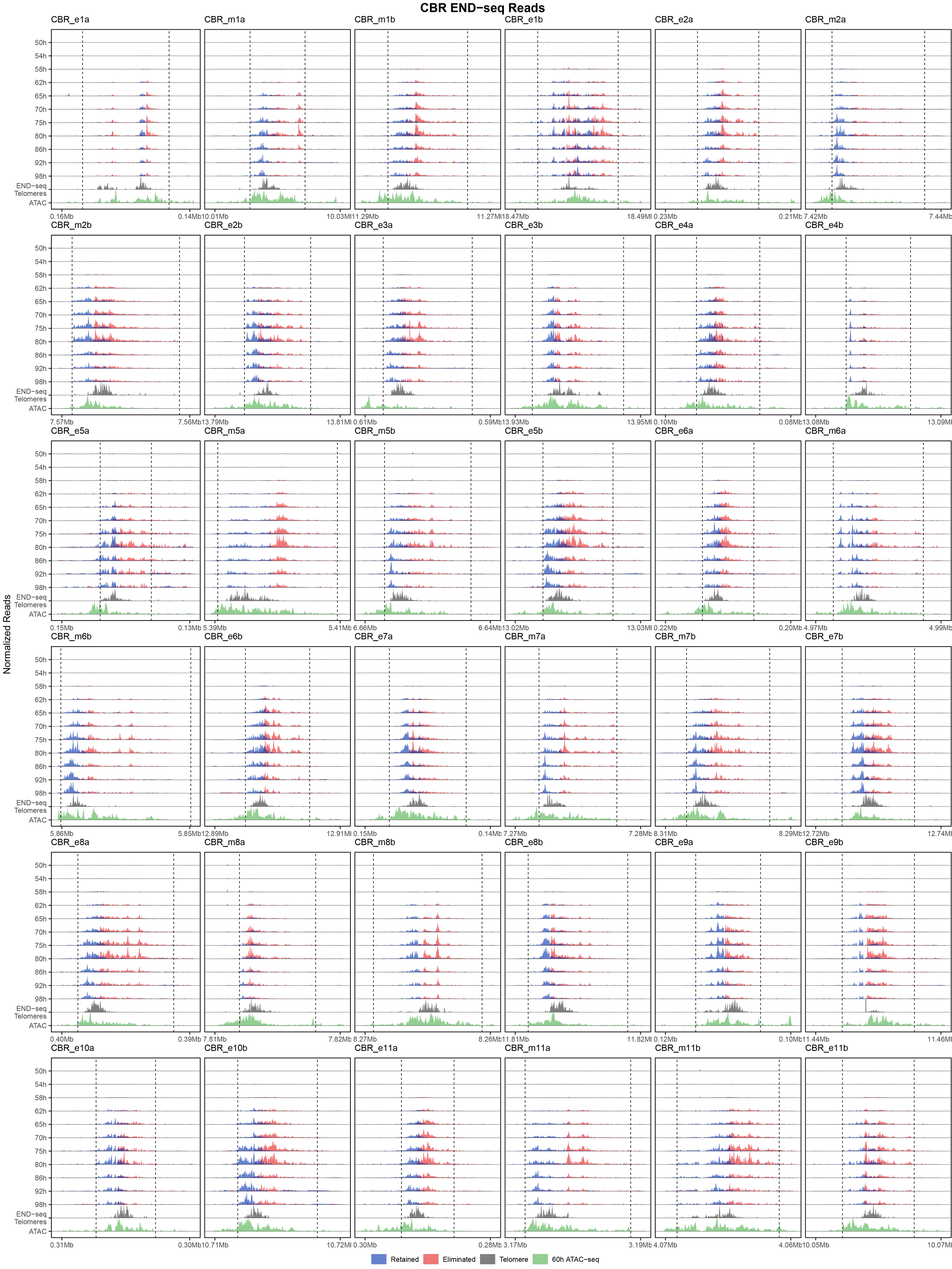
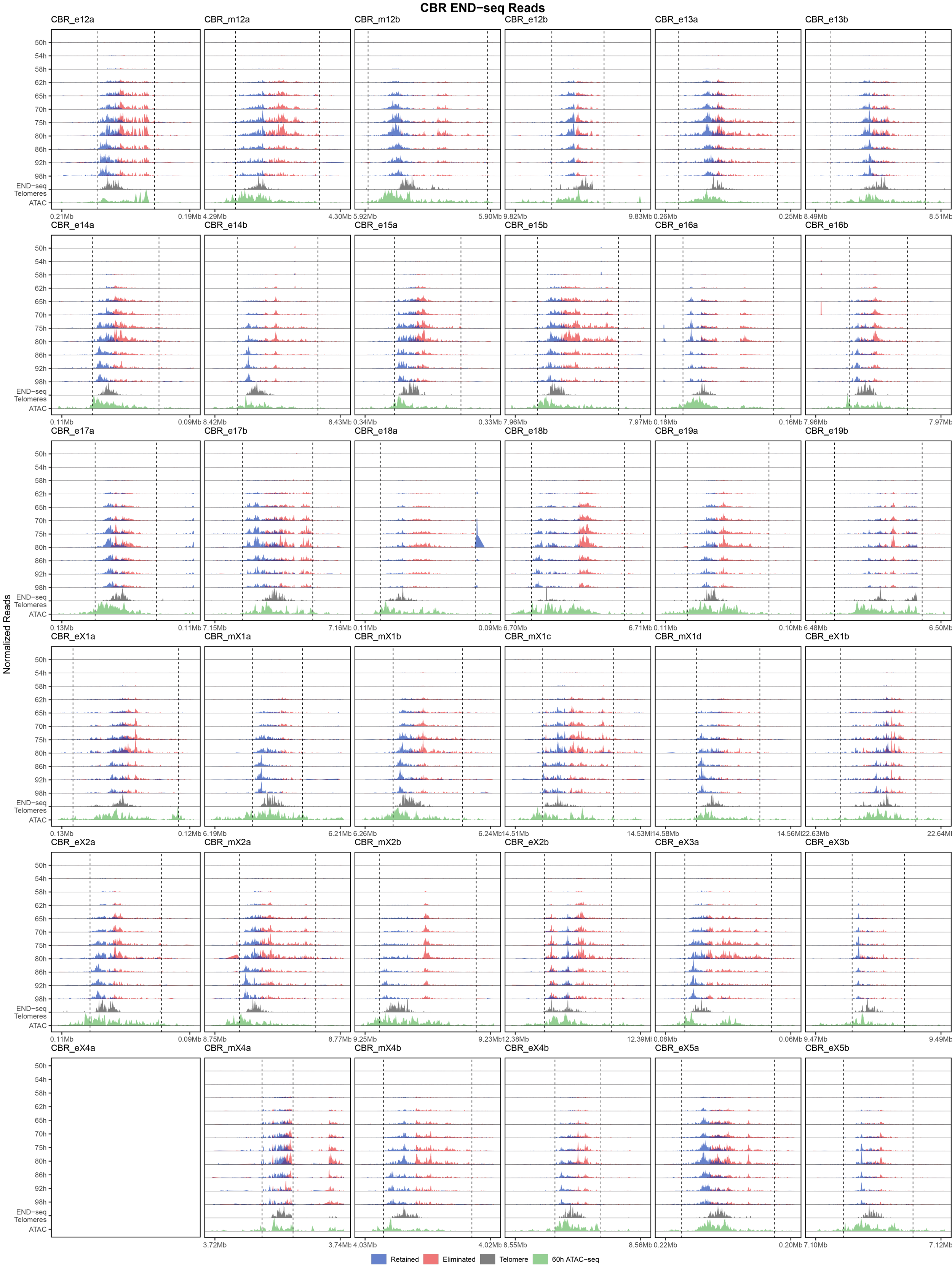
Timing of DSBs detected by END-seq. Ridgeline plots of END-seq reads across 11 developmental stages (y-axis) for each canonical CBR and its flanking regions (x-axis; total 20kb with 100 bp bins and 10 bp sliding window). Dashed lines mark the boundary of END-seq signal enrichment. Reads are colored by strand (red and blue), telomeric reads are grey, and ATAC-seq on 60h embryos is green. Plots are normalized to the highest END-seq signal at each CBR with telomeres and ATAC-seq independently normalized.

**Figure S2.**
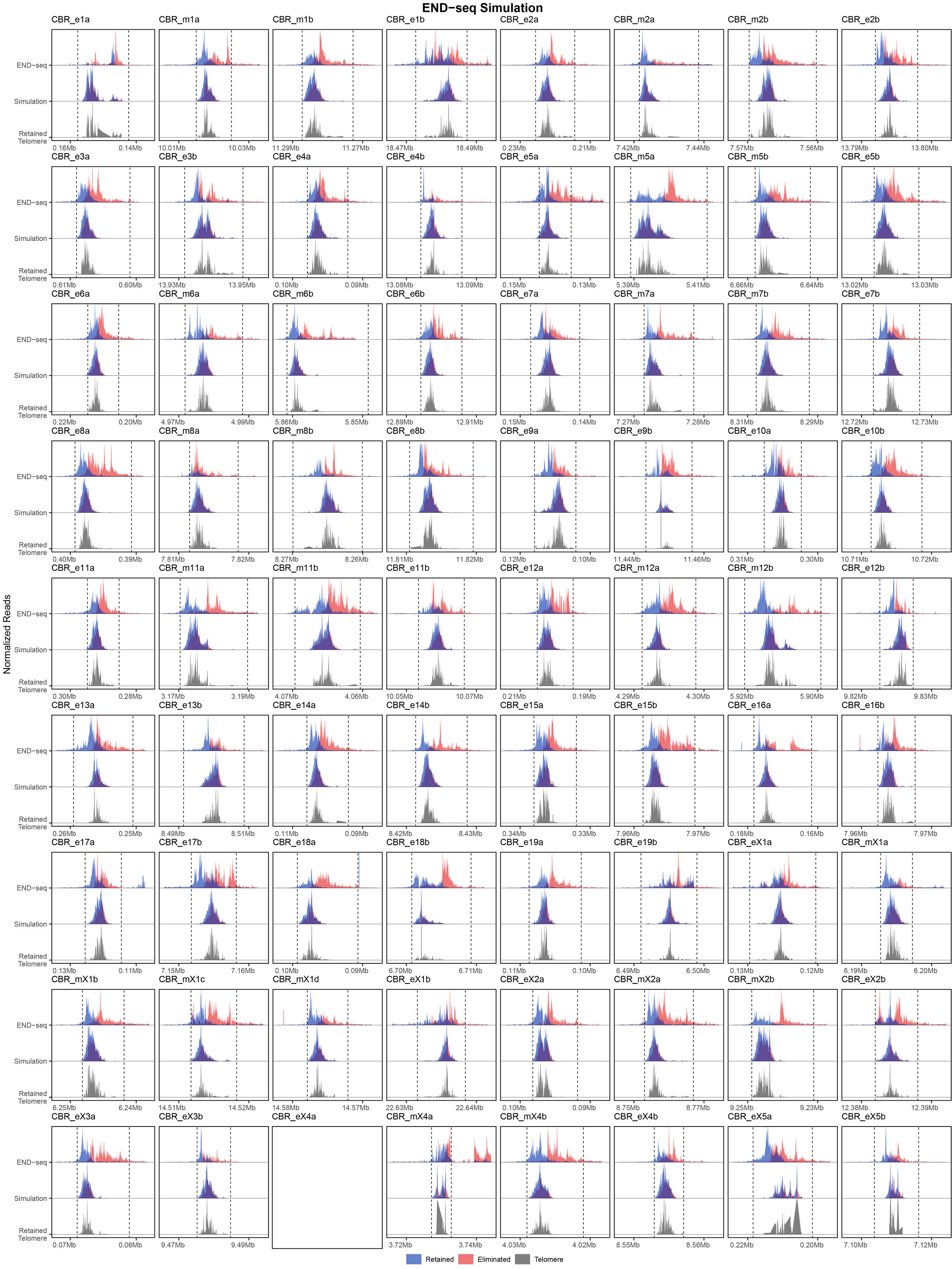
Simulation of END-seq pattern using telomere addition sites. Shown are observed *Ascaris* END-seq data (top) compared to simulated END-seq profiles (middle) using the END-seq profiles from *O. tipulae*, a nematode with homogeneous genetic background and homogeneous DSBs. The simulation used the position and frequency of *Ascaris* telomere addition sites (bottom) (see Methods). Each CBR is normalized independently to the highest read signal to better illustrate the differences within the CBRs.

**Table S1. CBRs and END-seq reads.** List of canonical and alternative CBRs and the number of END-seq reads in *Ascaris* and *Parascaris*.

**Table S2. CBR sequence conservation.** Sequence conservation was identified by pairwise BLAST comparison of CBRs and randomly sampled sequences within and between *Ascaris* and *Parascaris* (see methods).

**Table S3. Ascaris embryo development after X-ray.** The number of cells within an embryo was counted using light and fluorescence microscopy (Hoechst staining).

## AUTHOR CONTRIBUTIONS

B.E. and J.W. conceived the study. B.E., R.E.D., and J.W. carried out the investigation. B.E. and J.W. performed formal analysis. B.E. wrote the initial draft. B.E. and J.W. wrote the manuscript. R.E.D. and J.W. reviewed and edited the manuscript. J.W. provided supervision, project administration, and funding.

## ACKNOWLEDGEMENTS

We thank Bruce Bamber, Jeff Myers, and Routh Packing Co. for their support and hospitality in collecting *Ascaris* material; Martin Nielsen for the *Parascaris* material; Ryan Simmons for characterizing the x-ray irradiated embryos; and the University of Colorado Anschutz Medical Campus Genomics Core for sequencing services. We also thank Tom Dockendorff, Rachel Patton McCord, and Albrecht von Arnim for their comments and critical reading of the manuscript.

